# Prolonged fasting and glucocorticoid exposure drive dynamic DNA methylation in elephant seals

**DOI:** 10.1101/2025.05.23.654932

**Authors:** Emily F Gibson, Julia María Torres-Velarde, David C Ensminger, Diana D Moreno-Santillán, Daniel E Crocker, José Pablo Vázquez-Medina

**Author notes:** Correspondence: José Pablo Vázquez-Medina, Department of Integrative Biology, University of California, Berkeley, 3040 Valley Life Sciences Building #3140, Berkeley CA 94720-3140, or Julia María Torres-Velarde, Estancia posdoctoral por México, SECIHTI, Laboratorio de Biología Molecular y Cultivo Celular, Centro de Investigación en Alimentación y Desarrollo, Unidad Mazatlán, Av. Sábalo Cerritos, Cerritos, 82100 Mazatlán, Sinaloa, México. Equal contribution (Emily F Gibson, Julia María Torres-Velarde). Laboratorio de Biología Molecular y Cultivo Celular, Centro de Investigación en Alimentación y Desarrollo, Unidad Mazatlán (Julia María Torres-Velarde). Department of Biological Sciences, San Jose State University (David C Ensminger).

## Abstract

Elephant seals experience prolonged fasting while breeding, molting, and undergoing postnatal development. Fasting elephant seals adjust neuroendocrine function and gene expression to cope with potentially detrimental effects associated with extended fasting. DNA methylation alters gene expression by modulating accessibility to regions necessary to initiate transcription. The role of fasting and glucocorticoids on DNA methylation in elephant seals is understudied. We evaluated whether fasting alters global blood DNA methylation, the potential correlation between increased glucocorticoids and methylation, and the effects of glucocorticoids on DNA methylation in seal muscle cells in primary culture. We found that fasting transiently increases blood DNA methylation and that blood DNA methylation levels correlate with plasma cortisol. Hence, we then conducted bioinformatic analyses to identify regions in the elephant seal glucocorticoid receptor (GR) promoter that influence gene transcription through methylation (CPG islands). We identified one CpG island within the putative promoter region of GR gene. Methylation in this region, however, was unaffected by prolonged fasting. We then investigated whether exogenous glucocorticoids alter DNA methylation and gene expression profiles in seal muscle cells in primary culture (myotubes). Exposure to glucocorticoids for 12 or 48 hours decreased DNA methylation while upregulating pro-survival gene expression in seal myotubes. Our results show that whereas prolonged fasting transiently increases DNA methylation in elephant seal blood, sustained exposure to exogenous glucocorticoids decreases DNA methylation and activates a pro-survival transcriptional program in seal myotubes. Therefore, our results suggest that DNA methylation is a plastic, potentially cell-type-specific response that regulates gene expression in fasting seals.

**Summary statement:** DNA methylation is a plastic response to fasting and increased glucocorticoids in elephant seals. Our work underscores the role of epigenetic regulation of gene expression during energetically challenging conditions in seals and potentially other mammals.

## Introduction

Northern elephant seals (*Mirounga angustirostris)* spend most of their life feeding at sea, returning to land to breed and molt (Le Boeuf and Laws, 1994). While on land, elephant seals naturally experience prolonged periods of absolute food and water deprivation (fasting) (Champagne et al., 2012a). Elephant seal pups nurse for a month before fasting for two months while undergoing postnatal development (Le Boeuf and Laws, 1994). Prolonged fasting induces changes in gene expression and neuroendocrine function that likely help elephant seals cope with potentially detrimental effects associated with extended food and water deprivation, such as oxidant stress and inflammation (Ensminger et al., 2021; Torres-Velarde et al., 2021; Vázquez-Medina et al., 2010).

DNA methylation is a biochemical mechanism that regulates gene expression by reversibly adding a methyl (CH_3_) group to a nitrogenous base in the DNA sequence of promoter regions (Dudley et al., 2011; Jeltsch, 2006). CpG islands are promoters’ most common methylation areas (Bird, 1986; Gardiner-Garden and Frommer, 1987; Holliday and Pugh, 1975). Generally, DNA methylation suppresses gene expression due to the methyl groups interfering with protein-DNA and DNA-DNA interactions necessary to activate transcription (Boyes and Bird, 1992). Since DNA methylation is a reversible biological signal that can function as an on-off switch to control gene expression (Ramchandani et al., 1999), it represents a conserved strategy to quickly respond to changing environmental conditions.

Studies in true hibernators such as thirteen-lined ground squirrels and Syrian hamsters, torpid bats, and brumating Chinese alligators suggest that DNA methylation is an essential mechanism for regulating gene expression in response to changing environmental conditions (Alvarado et al., 2015; Coussement et al., 2023; Lin et al., 2020; Liu et al., 2023; Srere et al., 1992). However, the effects of fasting on DNA methylation in marine mammals have not been previously examined. In elephant seals, prolonged fasting increases glucocorticoid release (Ortiz et al., 2003; Ortiz et al., 2001), and previous work shows that glucocorticoid receptor (GR) signaling is crucial to elephant seal’s response to fasting (Avalos et al., 2023; Ensminger et al., 2021). In elephant seal muscle cells in primary culture, GR signaling induced by the synthetic glucocorticoid dexamethasone downregulates pro-inflammatory genes while upregulating genes that drive metabolic adjustments to support cell survival during energetic stress (Torres-Velarde et al., 2021).

Here, we studied global blood DNA methylation in fasting elephant seals and elephant seal muscle cells treated with exogenous glucocorticoids. We also analyzed the elephant seal GR promoter region for specific methylation patterns. We found that global blood DNA methylation increases reversibly with prolonged fasting, in concert with plasma cortisol levels in elephant seal pups. In contrast, sustained exposure to dexamethasone decreased DNA methylation while upregulating pro-survival genes in elephant seal muscle cells in primary culture. Hence, our results suggest that DNA methylation is a plastic, dynamic response that likely contributes to regulating gene expression in simultaneously fasting and developing seals. Our results also suggest that changes in DNA methylation in response to increased glucocorticoid treatment might be tissue- and gene-specific.

## Materials and methods

### Animal handling and sample collection

Animal work was approved by Sonoma State University and UC Berkeley Institutional Animal Care and Use Committees. Samples were collected under National Marine Fisheries Service Permit (NMFS) No. 1908 (PI Dan Costa, UCSC). Primary cells were derived under NMFS authorization No. 24479. Northern elephant seals were sampled during three time periods from February to October at Año Nuevo Reserve (San Mateo County, CA). The first sampling corresponded to early-fasting seal pups (< 1-week post-weaning, unmolted animals, male and female, N = 6). The second sampling corresponded to late-fasting pups (∼6 -weeks post-weaning, male and female fully molted animals, N = 6). The third sampling corresponded to post-foraging animals (∼8 months old, male and female, N =5) returning from their first oceanic feeding trip (Figure 1A). Animals were immobilized with tiletamine/zolazepam and ketamine as previously described (Vázquez-Medina et al., 2010). Blood samples were collected from the extradural vein into chilled EDTA vacutainers (BD Biosciences, San Jose, CA) and transported on ice to UC Berkeley. Plasma was separated by centrifugation, and the white blood cell layer was aliquoted and frozen at -80°C until DNA methylation analysis. Muscle biopsies were collected from the *longissimus dorsi* in the posterior flank of each animal using a 6 mm biopsy needle (Vázquez-Medina et al., 2010) from animals returning from their first oceanic feeding trip, rinsed with Hanks’ Balanced Salt Solution (Gibco, Waltham, MA), stored in cold Ham’s F10 nutrient mix (Gibco) supplemented with antibiotics, and transported on ice to UC Berkeley for cell isolation (Torres-Velarde et al., 2021).

**Figure 1.**
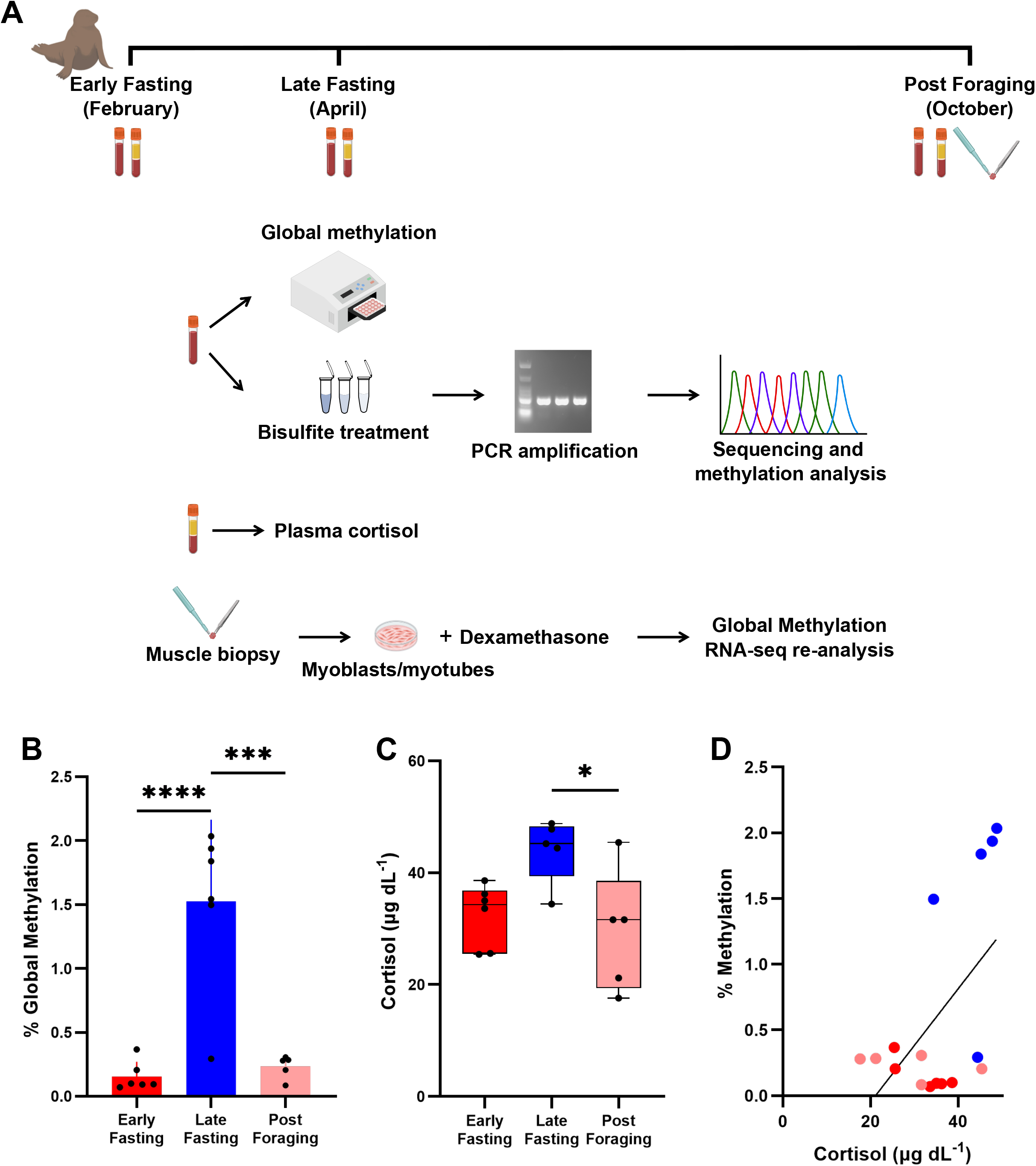
Fasting increases blood DNA methylation in elephant seals. **A)** Sampling protocol. **B)** Blood DNA methylation in elephant seals during early, late, and post-foraging periods. **C)** Plasma cortisol concentrations in early, late, and post-foraging animals. * = *p<0*.*05*. Data are mean ± SD. **D)** Correlation of plasma cortisol and global blood DNA methylation; Deming linear regression model: Y = 13.82*X - 4.25, 95% confidence interval intercept -7.429 to - 1.075, slope 5.674 to 21.97 (r = 0.554, *p=0*.*0255*). Dots represent individual animals.

### DNA extraction, global DNA methylation, and cortisol measurements

Genomic DNA was extracted using the salting method (Miller et al., 1988). Samples were incubated with lysis buffer (5M NaCl, 1M Tris, 0.5M EDTA, 10% sodium dodecyl sulfate, Invitrogen, Waltham, MA) and proteinase K (Promega, Madison, WI) overnight at 55°C. DNA was precipitated with 100% ethanol, washed twice with 70% ethanol, and resuspended in Tris-EDTA (Invitrogen). DNA was purified using a Wizard^®^ SV Gel and PCR Clean-Up System kit (Promega) and quantified using a NanoDrop spectrophotometer (Nanodrop Technologies, Wilmington, DE) and a Qubit fluorometer (Invitrogen). Percent global DNA methylation was quantified in triplicate (CV=15.2) using 100 ng of DNA and a colorimetric Global DNA Methylation Assay kit (Abcam, Cambridge, UK). Plasma cortisol (CV=6.9) was quantified in 20 µl of plasma diluted 1:2 using an enzyme-linked immunosorbent assay (Alpco, Salem, NH) previously validated for elephant seals (McCormley et al., 2018).

### Promoter identification and characterization

The Northern elephant seal glucocorticoid receptor (GR) gene was identified based on a sequence similarity search by comparing the database of annotated sequences in the elephant seal genome (Torres-Velarde et al., 2021) to the human GR (UniProt: P04150). The similarity search was conducted using DIAMOND (double index alignment of next-generation sequencing data) under the “more-sensitive” parameter (Buchfink et al., 2015). Corresponding promoter sequences were identified upstream of the elephant seal GR gene by aligning the elephant seal sequences with publicly available marine mammal sequences using BLAST (https://blast.ncbi.nlm.nih.gov). Once the putative promoter region was identified, 3,341 base pairs upstream of the start codon of GR, including the 5’ untranslated region, were used to identify CpG islands with MethPrimer ((Li and Dahiya, 2002), http://www.urogene.org/cgi-bin/methprimer/methprimer.cgi). Transcription factor binding sites were identified within the CpG islands using Nsite (http://www.softberry.com/berry.phtml?topic=nsite&group=programs&subgroup=promoter). The Transcription Start Site (TSS) was identified using TSSG software (Shahmuradov and Solovyev, 2015; Solovyev et al., 2010). Only regions with 0 mismatch between the elephant seal genome and the transcription factor binding site were considered for further analysis.

### Bisulfite conversion, PCR amplification, Sanger sequencing, and methylation analysis

One µg of genomic DNA was converted with sodium bisulfite using the Qiagen EpiTect Bisulfite Kit (Qiagen, Hilden, Germany). The concentration of bisulfite-converted DNA was measured using a Nanodrop spectrophotometer (Nanodrop Technologies). Bisulfite-sequencing PCR (BSP) primers were designed using the MethPrimer tool (Li and Dahiya, 2002, https://www.methprimer.com/cgi-bin/methprimer/methprimer.cgi) to amplify fragments of about 300 bp and synthesized by Integrated DNA Technologies (Coralville, IA). The GR BSP1 primers were designed to amplify a region in the CpG island 259 base pairs long, which contained 56 CpG sites. The GR BSP2 primers were designed to amplify a region in the CpG island 339 base pairs long, with 66 CpG sites. The GR BSP3 primers were designed to amplify a region in the CpG island about 288 base pairs long, which contained 28 CpG sites. Primer sequences are listed in Table 1. PCR fragments were amplified using Platinum Taq DNA Polymerase (Thermo Fisher Scientific, Waltham, MA), purified using the Wizard^®^ SV Gel and PCR Clean-Up System (Promega), and verified on 2% agarose gels. Amplified fragments were Sanger-sequenced at the UC Berkeley DNA sequencing facility. Sequences were analyzed using the Quantification tool for Methylation Analysis (QUMA, http://quma.cdb.riken.jp/) to evaluate the bisulfite conversion rate and determine CpG methylation site frequency. Sequences with less than 90% conversion rates were excluded from the analysis.

**Table 1:**
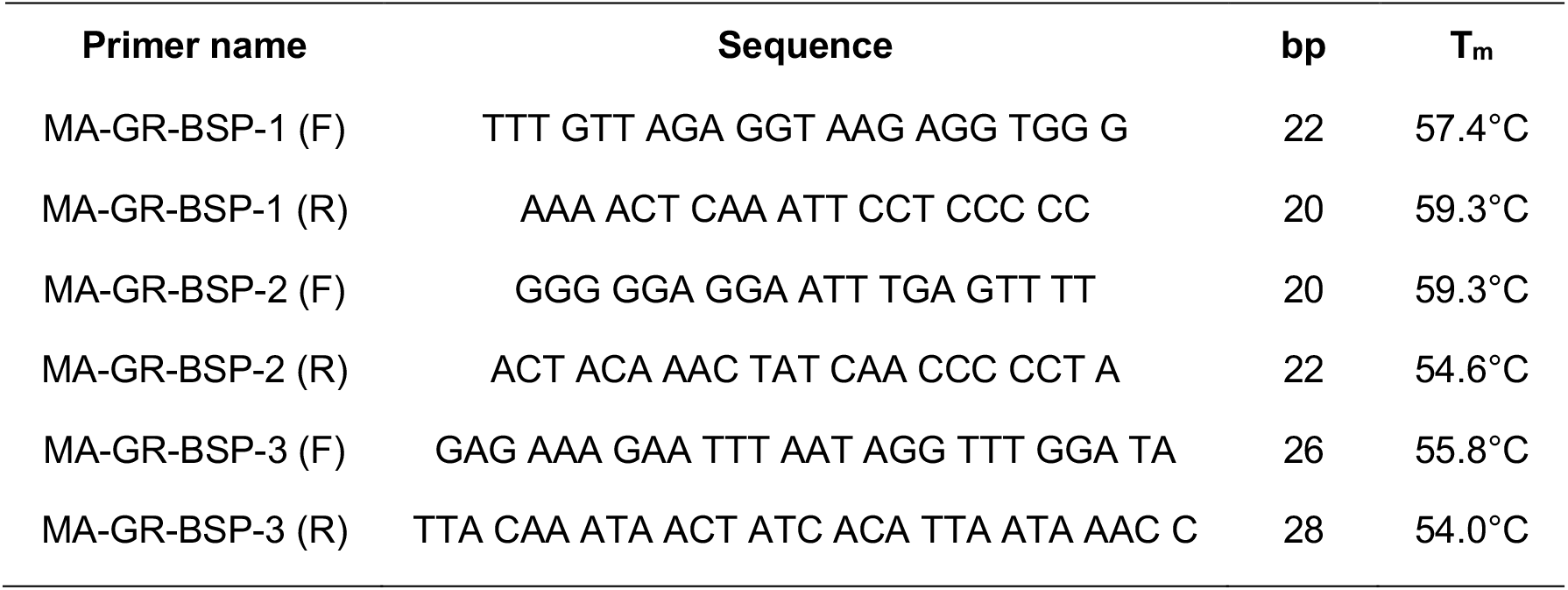
Primers used to amplify and sequence regions in the CPG island of the elephant seal GR promoter.

### Primary cell isolation, differentiation, and dexamethasone treatment

Muscle progenitor cells were isolated from muscle biopsies and characterized as described in our previous work (Torres-Velarde et al., 2021). Cells derived from three post-foraging animals were seeded in tissue culture dishes coated with 0.01% collagen (Sigma, St. Louis, MO) in Ham’s F10 nutrient mix (Gibco) supplemented with 20% fetal bovine serum (Seradigm, Radnor, PA), 100mM HEPES (Gibco), and 1% Antibiotic-Antimycotic solution (Gibco). Myotube differentiation was induced by incubation in DMEM (Gibco) supplemented with 5% horse serum (Gibco) for seven days. This protocol produces highly differentiated myotubes with distinctive gene expression profiles, morphological and metabolic phenotypes (Torres-Velarde et al., 2021). Myotube differentiation was confirmed by microscopy (Figure 3A). Myotubes were treated with the synthetic glucocorticoid dexamethasone (BioVision, Milpitas, CA) at three ascending concentrations (0.1µM, 1.0µM, and 100µM) for 6, 12, or 48h. Dexamethasone induces a prolonged GR-mediated response while minimizing activation of the mineralocorticoid receptor in various cells, including elephant seal myotubes (Torres-Velarde et al., 2021). DMSO was used as a vehicle control. Cells were collected for DNA extraction after treatment and analyzed for global DNA methylation using a colorimetric Global DNA Methylation Assay kit (Abcam).

### Re-analysis of previously generated RNA-seq data

RNA-seq data from elephant seal myotubes treated with 100µM dexamethasone for 48h (Torres-Velarde et al., 2021) were reanalyzed using our previously published pipeline with minor modifications (Allen et al., 2024; Torres-Velarde et al., 2021). Total RNA was extracted using TRIzol (Invitrogen, Carlsbad, CA). RNA integrity was measured using a Bioanalyzer (Agilent Technologies, Santa Clara, CA). RNA with RIN > 9.5 was used to prepare cDNA libraries from poly(A)-captured mRNA using a KAPA RNA HyperPrep Kit (Roche, Basel, Switzerland, Cat. No. KK8541) with TruSeq adapters from three replicates per treatment. A total of 25 M reads per sample were sequenced on a NovaSeq platform (Illumina, San Diego, CA) at the UC Berkeley Functional Genomics and Vincent J. Coates Genomics Sequencing Laboratories. Reads were mapped to the elephant seal genome (https://www.dnazoo.org/assemblies/mirounga_angustirostris) annotated in (Torres-Velarde et al., 2021). Unannotated genes were manually curated and annotated using the latest version of the human genome reference (Ensembl GRCh38.p14). A BLAST search was performed against the human genome to identify homologous sequences, and Ensembl gene IDs were converted to gene symbols using the BioMart tool. Transcript levels were quantified using RSEM 1.3.1 (Li and Dewey, 2011). Genes differentially expressed (DE) between control and 100µM dexamethasone were identified at an FDR of 5% using EBSeq (Leng et al., 2013). Over-representation analysis (ORA) with gene ontology (GO) biological processes, KEEG, and Reactome databases were conducted using Webgestalt (Elizarraras et al., 2024) to identify enriched pathways in upregulated genes (FDR <0.05). Cis-regulatory element analysis was performed using iRegulon in Cytoscape at an FDR < 0.001 to identify the top 3 transcription factors acting on upregulated genes (Janky et al., 2014).

### Statistical analyses

All analyses were conducted in GraphPad Prism v10.1.1. Normality and homoscedasticity were assessed using D’Agostino-Pearson and Kolmogorov-Smirnov tests. Group differences were evaluated using ANOVA or Kruskal-Wallis tests, with Fisher’s or Dunn’s post hoc corrections, based on the mean of triplicate measurements per biological sample (individual seal or cell line). The relationship between global DNA methylation and cortisol was examined using Pearson correlation and Deming regression linear models. Differences were considered significant at p<0.05. Data are presented as mean ± SD.

## Results

### Prolonged fasting induces reversible blood DNA methylation in elephant seals

We measured the effects of prolonged fasting on plasma cortisol and global blood DNA methylation in elephant seals. Mean methylation values were ten times higher in late than early fasting seals and returned to early fasting levels in post-foraging animals (KW = 10.24, *p=0*.*0015*, Figure 1B, Supplementary table 1). These data show that prolonged fasting alters DNA methylation profiles transiently in elephant seal blood. Whereas plasma cortisol levels were not statistically higher in late than in early fasting seals (F = 5.206, *p=0*.*0608*), they were significantly lower in post-foraging than in late fasting animals (F = 5.206, *p=0*.*0246*) (Figure 1C). Moreover, we detected a statistically significant positive correlation between plasma cortisol and global blood DNA methylation levels (*r=0*.*554, p=0*.*025*5), suggesting that blood DNA methylation increases with cortisol throughout the fasting period (Figure 1D).

### The putative promoter region in the elephant seal GR gene contains one CpG island, which is not methylated in response to prolonged fasting in elephant seal blood

We searched for CpG islands in the elephant seal GR gene and looked for changes in GR methylation in this region in response to fasting in elephant seal blood. We detected one CpG island within the 3,338 base pair region of the elephant seal GR promoter (Figure 2A). This CpG island is 1,747 bp long, located between -1909 nt and -163 nt, upstream of the transcription start site (Figure 2A). We then annotated transcription factor binding motifs and other regulatory elements in the sequenced region of the elephant seal GR promoter (Figure 2B). The region of the GR promoter amplified by the BSP1 primer pair contained five regulatory element binding motifs, including binding sites for Oct-1-2-4, TGT3, cyclin D2, ETF-A, USF1, 2, PU.1, and GABPα transcription factors. The GR promoter region amplified by the BSP2 primer pair contained three regulatory element binding motifs, including sites for E2F, SP1-3, and HIF-1 transcription factors. The region of the GR promoter amplified by the BSP3 primer pair contained four regulatory element binding motifs, including sites for Smad3-4, SP1, E4BP4, and AP2 transcription factors (Figure 2B). No CpG sites were methylated in any of the blood samples (early fasting, late fasting, or post-foraging) (Supplementary Figures 1, 2, 3). These data show that in this region of the GR promoter, methylation does not change in response to prolonged fasting in blood cells, suggesting that if other transcription factors regulate the expression of GR or other nearby genes, such regulation is likely not affected by methylation, at least in blood cells.

**Figure 2.**
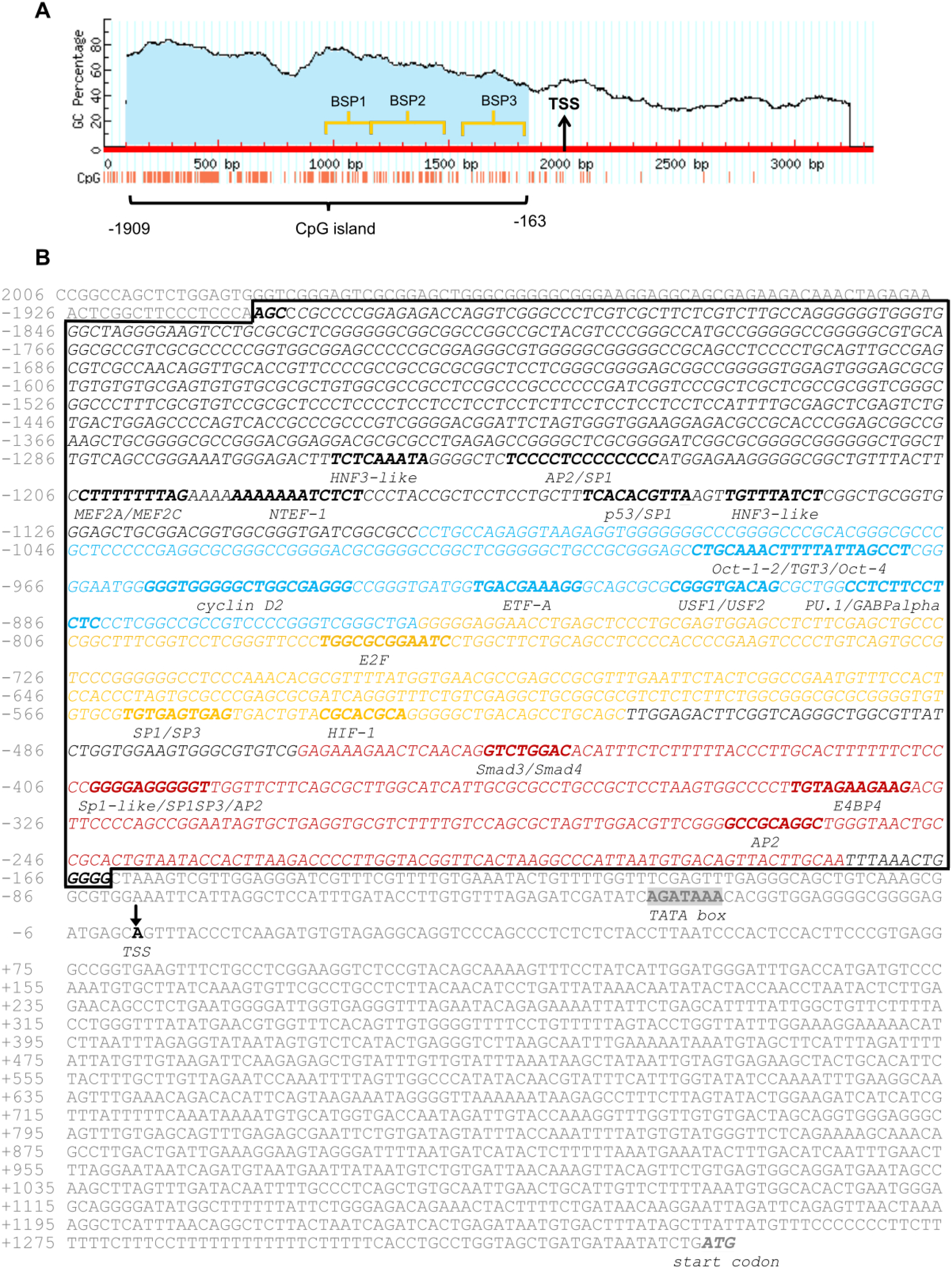
A) Identification of the CpG island (light, blue-shaded area) in the putative promoter region of the elephant seal GR gene. CpG was identified between -1908 and -162 nt position (1747 bp length) upstream of the TSS. **B)** Annotation of possible transcription factor binding motifs and other regulatory elements in the sequenced region of the elephant seal GR promoter (black and italic). Nucleotides in blue correspond to the section of DNA amplified by the primer pair MA-GR-BSP1; sequences in yellow correspond to the section of DNA amplified by the primer pair MA-GR-BSP2; sequences in red correspond to the section of DNA amplified by the primer pair MA-GR-BSP3. Primers were designed to amplify specific regions of the CpG island in the elephant seal GR promoter for methylation status characterization. The black box indicates the CpG island region. TATA box (black and gray), Transcription start site (TSS) (black arrow) and start codon (ATG) (black and gray). Analyses were conducted using Softberry NSITE software (http://www.softberry.com/berry.phtml?topic=nsite&group=programs&subgroup=promoter&advanced=on)

**Figure 3.**
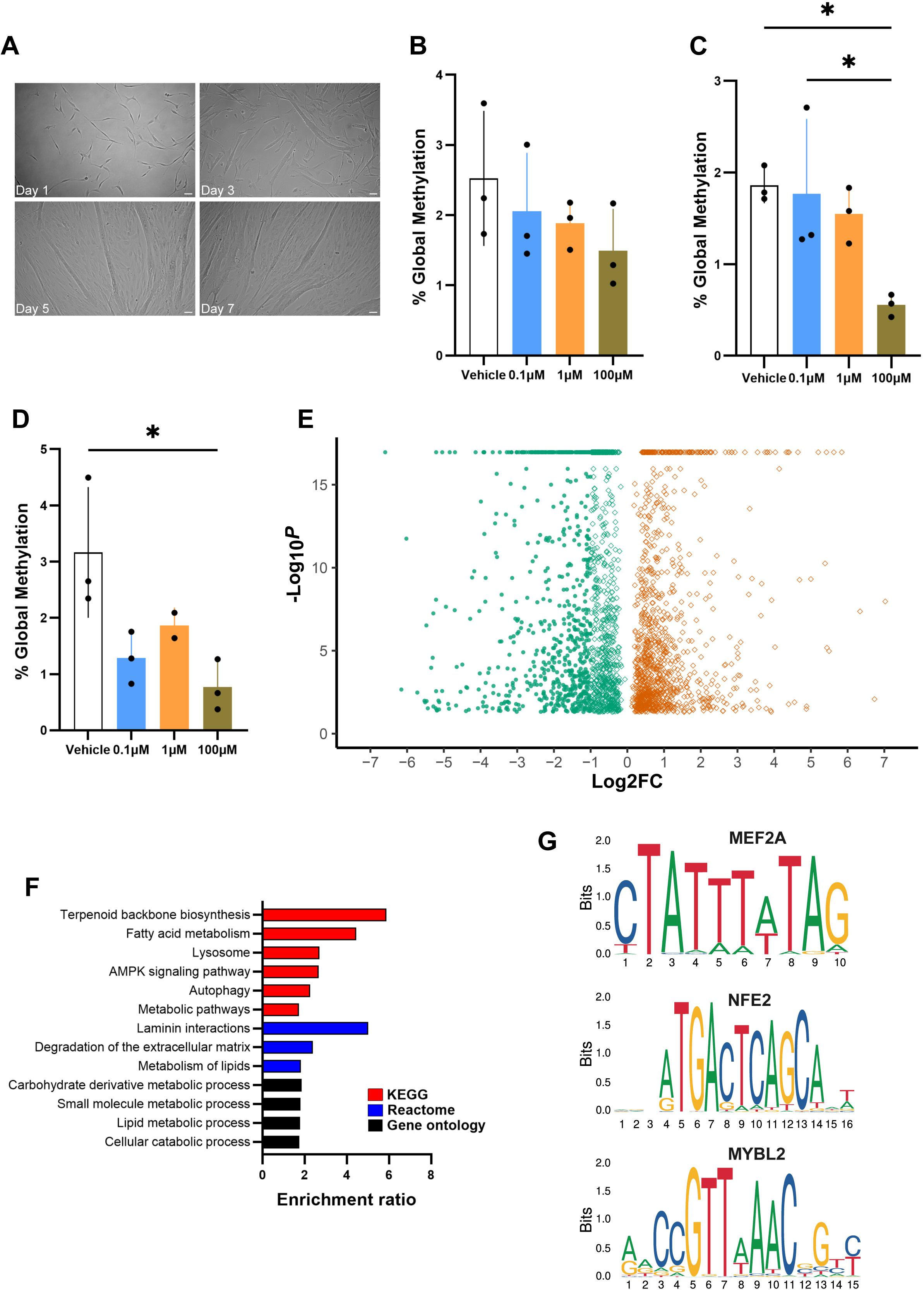
Treatment with glucocorticoids decreases global methylation in elephant seal myotubes. **A)** Elephant seal myoblasts differentiating into myotubes. DNA methylation in cells treated with ascending concentrations of dexamethasone (0.1, 1.0, and 100µM) for **B)** 6, **C)** 12, or **D)** 48 hours. * indicates *p<0*.*05*. Data are mean ± SD. **E)** Volcano plot showing differentially expressed genes in elephant seal myotubes treated with 100µM dexamethasone for 48h (FDR <0.05). **F)** Over-representation analyses (ORA) in upregulated DE genes (FDR <0.05). **G)** Top enriched motifs identified by cis-regulatory element analysis of RNA-seq data. Sequence logos were retrieved from the Jaspar 2020 database (https://jaspar2020.genereg.net/).

### Treatment with synthetic glucocorticoids decreases DNA methylation while upregulating metabolic and pro-survival genes in elephant seal myotubes

We tested whether exogenous treatment with the synthetic glucocorticoid dexamethasone, which induces potent, GR-specific signaling, alters global DNA methylation in elephant seal myotubes in primary culture. Treatment with ascending dexamethasone concentrations for 12 and 48, but not 6 hours, decreased global methylation only at the highest dose (100µM, Figures 3B, C, D). These data suggest that elephant seal myotubes retain the capacity to respond to sustained high levels of glucocorticoids, potentially by regulating gene transcription. Hence, we re-analyzed RNA-seq data previously generated from elephant seal myotubes treated with 100µM dexamethasone for 48h. As described in our previous work (Torres-Velarde et al., 2021), 2,488 genes were differentially expressed (FDR < 0.05%) in cells treated with dexamethasone compared to control, including 1,078 upregulated genes and 1,410 downregulated genes (Figure 3E). 2,406 of the 2,488 differentially expressed genes were annotated, including 1,045 upregulated genes and 1,361 downregulated genes. Notably, dexamethasone treatment decreased GR expression (NR3C1) by 60% (not shown), further suggesting that elephant seal GR expression is not regulated by methylation. DNA methylation generally suppresses gene expression (Boyle et al., 2008; Newell-Price et al., 2000), and our experiments show that dexamethasone treatment decreases DNA methylation in elephant seal myotubes. Therefore, we focused on differentially expressed genes upregulated by dexamethasone. Overrepresentation analyses for GO biological processes, KEEG, and Reactome pathways showed strong signatures for genes involved in muscle remodeling, energy metabolism, and cell survival (Figure 3F). We then analyzed cis-regulatory elements to identify enriched motifs for transcription factors acting on upregulated genes. Myocyte-specific enhancer factor 2A (MEF2A), Nuclear factor, erythroid 2 (NFE2), and Myb-related protein B (MYBL2) were the top 3 predicted transcription factors with normalized enriched scores of 5.069 (MEF2A), 5.054 (NFE2), and 4.618 (MYBL2) (Figure 3G). MEF2 regulates skeletal muscle cell differentiation (Black and Olson, 1998; Edmondson et al., 1994) and cell survival via p38 MAPK signaling and interaction with STAT transcription factors (Moustafa et al., 2023). NFE2 is a known regulator of skeletal muscle (Llano-Diez et al., 2011), which interacts with CREB-binding protein (Yao et al., 2018), regulating transcription factors and genes responsible for lipogenesis and gluconeogenesis (Hung et al., 2001). NEF2 is also crucial to regulating antioxidant gene expression through AP1 binding sites (Shyu et al., 2005). MYBL2 regulates cell cycle progression, survival, and differentiation via transcriptional regulation and direct interaction with serine-threonine kinase receptor-associated protein (STRAP), which suppresses TGF-β signaling (Musa et al., 2017). Hence, our data suggest that the observed decrease in DNA methylation in seal myotubes treated with dexamethasone is associated with a pro-survival transcriptional response that helps these cells cope with sustained glucocorticoid elevations (Torres-Velarde et al., 2021).

## Discussion

DNA methylation is an adaptive mechanism that regulates gene expression in response to environmental stimuli (Reik, 2007). Here, we show that prolonged fasting transiently increases genome-wide blood DNA methylation in elephant seals in parallel with increasing plasma cortisol. Interestingly, our results also show that sustained exposure to the synthetic glucocorticoid dexamethasone decreases DNA methylation in elephant seal muscle cells in primary culture while upregulating the expression of metabolic and pro-survival genes. Hence, DNA methylation is potentially a cell-type-specific strategy to fine-tune gene expression during energetically challenging conditions in elephant seals. Previous work showed that fasting elephant seals experience substantial systemic physiological adjustments. However, such adjustments are largely tissue-specific, especially in seal pups, which simultaneously undergo fasting and postnatal development (Viscarra et al., 2013).

Previous studies also show that cyclical patterns in DNA methylation are a transient response to environmental stress (Alvarado et al., 2015; Coussement et al., 2023; Kangaspeska et al., 2008; Lin et al., 2020; Secco et al., 2015; Srere et al., 1992), supporting our conclusion that fasting-induced blood DNA methylation in elephant seals is a plastic mechanism to modify gene expression in response to environmental cues. Along with epigenetic changes in white blood cells, we detected changes in circulating cortisol in fasting seals. Post-weaning fasting increases plasma cortisol in elephant seal pups (Ortiz et al., 2003; Ortiz et al., 2001; Viscarra et al., 2011), likely contributing to lipid oxidation and endogenous glucose production (Champagne et al., 2013; Champagne et al., 2005; Champagne et al., 2012b; Rea and Costa, 1992; Viscarra et al., 2012). Hence, our results suggest that concerted systemic changes in plasma cortisol and blood DNA methylation may contribute to sustaining metabolic functions in fasting elephant seals.

While measuring global methylation shows overall changes over time, it does not provide details on specific methylated genes. Hence, we used bisulfite conversion, PCR amplification, and Sanger sequencing (Zhang et al., 2009) to construct a specific methylation profile for the GR promoter, which regulates the neuroendocrine stress response to prolonged fasting and mediates metabolic adaptation in elephant seal muscle cells (Ensminger et al., 2021; Torres-Velarde et al., 2021). We sequenced regions within the CPG island of the GR promoter with three sets of BSP primers but did not find changes in blood GR methylation in fasting seal blood. Moreover, we found that while sustained exposure to exogenous glucocorticoids decreases DNA methylation and upregulates pro-survival genes in elephant seal muscle cells, it suppresses GR expression, further suggesting that either GR is not regulated by methylation or that a negative feedback mechanism suppresses GR expression during glucocorticoid elevations (Torres-Velarde et al., 2021). Of note, we did not evaluate other epigenetic mechanisms that might be involved in regulating GR expression, such as histone modifications (McCaw et al., 2020; Turner, 2002) or microRNAs (De Falco et al., 2023; Penso-Dolfin et al., 2020). Furthermore, while treatment with exogenous synthetic glucocorticoids is a powerful tool to induce GR signaling in cultured cells, it might not accurately recapitulate fasting-induced increases in circulating cortisol. Hence, more *in vivo* studies that directly evaluate epigenetic changes in complex tissues under natural fasting conditions are needed.

Analysis of the sequenced region of the elephant seal GR promoter identified several binding sites for transcription factors that regulate chromatin remodeling (Kuznetsova et al., 2015; McDowell et al., 2018), cellular proliferation (Gay et al., 2016), the immune response (Adcock, 2001), and oxidant stress (You et al., 2009). Elephant seals are adapted to cope with oxidant stress induced by prolonged fasting and extended breath-hold diving (Crocker et al., 2016; Vázquez-Medina et al., 2012). GR can interact with AP1 (Vesely et al., 2009), regulating the transcriptional response to oxidant stress (Zhou et al., 2001). We identified multiple AP1 binding sites within the sequenced region of the GR promoter, indicating that more than one AP1 subunit is involved in regulating GR expression. Our bisulfite sequencing results, however, showed that no binding sites were methylated in fasting elephant seals’ blood cells. Furthermore, cis-regulatory element analysis of genes upregulated in elephant seal muscle cells treated with exogenous synthetic glucocorticoids identified NEF2, which regulates antioxidant gene expression through AP1 binding sites (Shyu et al., 2005), as an enriched transcription factor. AP1 has both positive and negative interactions with GR (Biddie et al., 2011; Yang-Yen et al., 1990), and is often activated in response to oxidant stress, but can also interact with GR during non-stress-mediated chromatin remodeling and mutually inhibitory protein-protein interactions. The lack of methylation in AP1 binding motifs despite the observed increases in global DNA methylation in seal blood suggests that GR and AP1 interact, potentially regulating antioxidant gene transcription during prolonged fasting (Crocker et al., 2016; Vázquez-Medina et al., 2010; Vázquez-Medina et al., 2013; Vázquez-Medina et al., 2011).

DNA methylation is highly cell-type-specific (Brandeis et al., 1993; Eden and Cedar, 1994) and plastic, with methylation or demethylation patterns regulating cellular differentiation throughout development (Carrió and Suelves, 2015). We observed opposite methylation patterns in white blood cells from fasting animals and cultured myotubes treated with exogenous synthetic glucocorticoids. During fasting, elephant seal weaned pups maintain muscle development without experiencing muscle atrophy, and elephant seal muscle cells use alternative metabolic pathways to support energy metabolism during sustained exposure to synthetic glucocorticoids (Torres-Velarde et al., 2021; Wright et al., 2020). The observed decrease in global methylation in response to sustained dexamethasone exposure in muscle cells and the over-representation of metabolic pathways in upregulated genes suggest that methylation might be an important mechanism to regulate genes involved in muscle development. Previous studies posited that DNA demethylation increases the expression of muscle proliferation genes (Brunk et al., 1996; Chiu and Blau, 1985). Further in vivo work is needed to understand how changes in methylation regulate gene expression in skeletal muscle at different time points during fasting.

In summary, we found that prolonged fasting increases global DNA methylation transiently in elephant seal blood, that blood DNA methylation profiles correlate with plasma cortisol levels, and that treatment with synthetic glucocorticoids decreases DNA methylation while upregulating pro-survival and metabolic genes in elephant seal muscle cells in primary culture. Hence, DNA methylation is a dynamic, potentially cell-type-specific response that modulates gene expression in elephant seals. Future work on tissue- and gene-specific changes in DNA methylation patterns across the entire genome would contribute to our understanding of DNA methylation’s dynamic role in regulating gene expression in fasting seals. Using next-generation sequencing, such as whole-genome bisulfite sequencing (WGBS), would provide a more comprehensive view of the interactions between DNA methylation patterns, environmental factors, and epigenetic modifications, increasing our understanding of gene expression regulation in fasting seals.

## List of symbols and abbreviations

GR: Glucocorticoid receptor
EDTA: Ethylenediaminetetraacetic acid
HEPES: N-2-hydroxyethyl piperazine-N-2-ethane sulfonic acid
DMEM: Dulbecco’s Modified Eagle Medium
DMSO: Dimethyl sulfoxide
PCR: Polymerase chain reaction
DIAMOND: Double index alignment of next-generation sequencing data
BSP: Bisulfite-sequencing PCR
QUMA: Quantification tool for methylation analysis

## Acknowledgments

EFG was supported by the UC Berkeley Undergraduate Research Apprenticeship Program (URAP) and a Chandra research fellowship through the Berkeley Summer Undergraduate Research Fellowship (SURF) program. JMT-V was supported by a UC MEXUS-CONACYT postdoctoral fellowship. DCE was supported by an NSF PRFB. Analysis of fragmented DNA was conducted at the UC Berkeley Functional Genomics Lab. Bisulfite sequencing was conducted at UC Berkeley’s DNA Sequencing facility. We thank Kaitlin Allen and Emily Lam for their assistance in the field and Peter Sudmant and Rohit Kolora for annotating the elephant seal genome.

## Data and resource availability

All relevant data and resources can be found within the article and its supplementary information.

---

**Figure S1.**
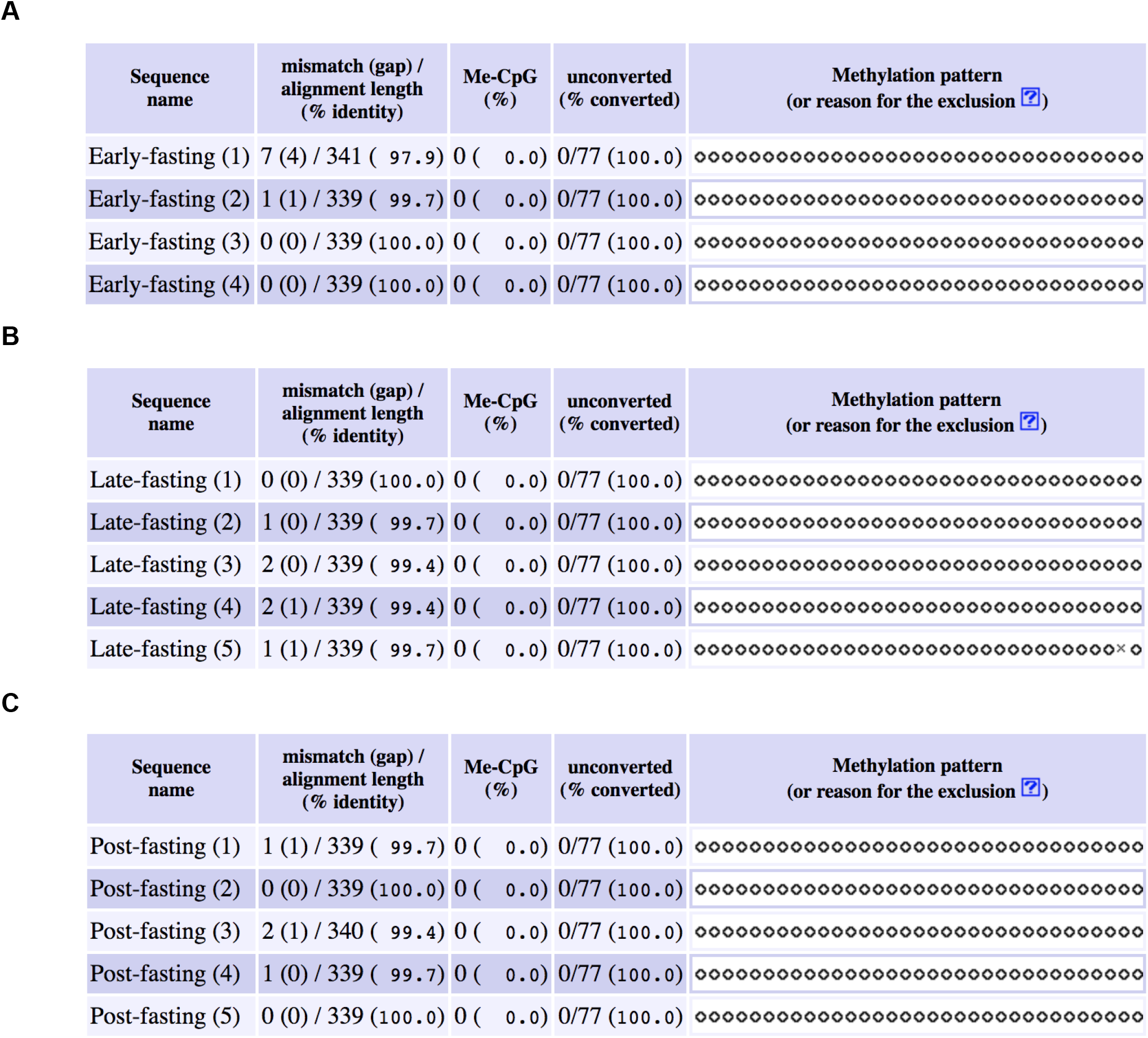
Methylation status in the GR promoter region (based on sequencing using BSP1 primers) in A) early fasting, B) late fasting, and C) post-foraging elephant seal pups. Filled-in (dark) circles indicate methylated CpG sites, and unfilled circles indicate unmethylated CpG sites. An “x” instead of a circle indicates that the CpG site was not sequenced. Data was compiled using QUMA software (http://quma.cdb.riken.jp/).

**Figure S2.**
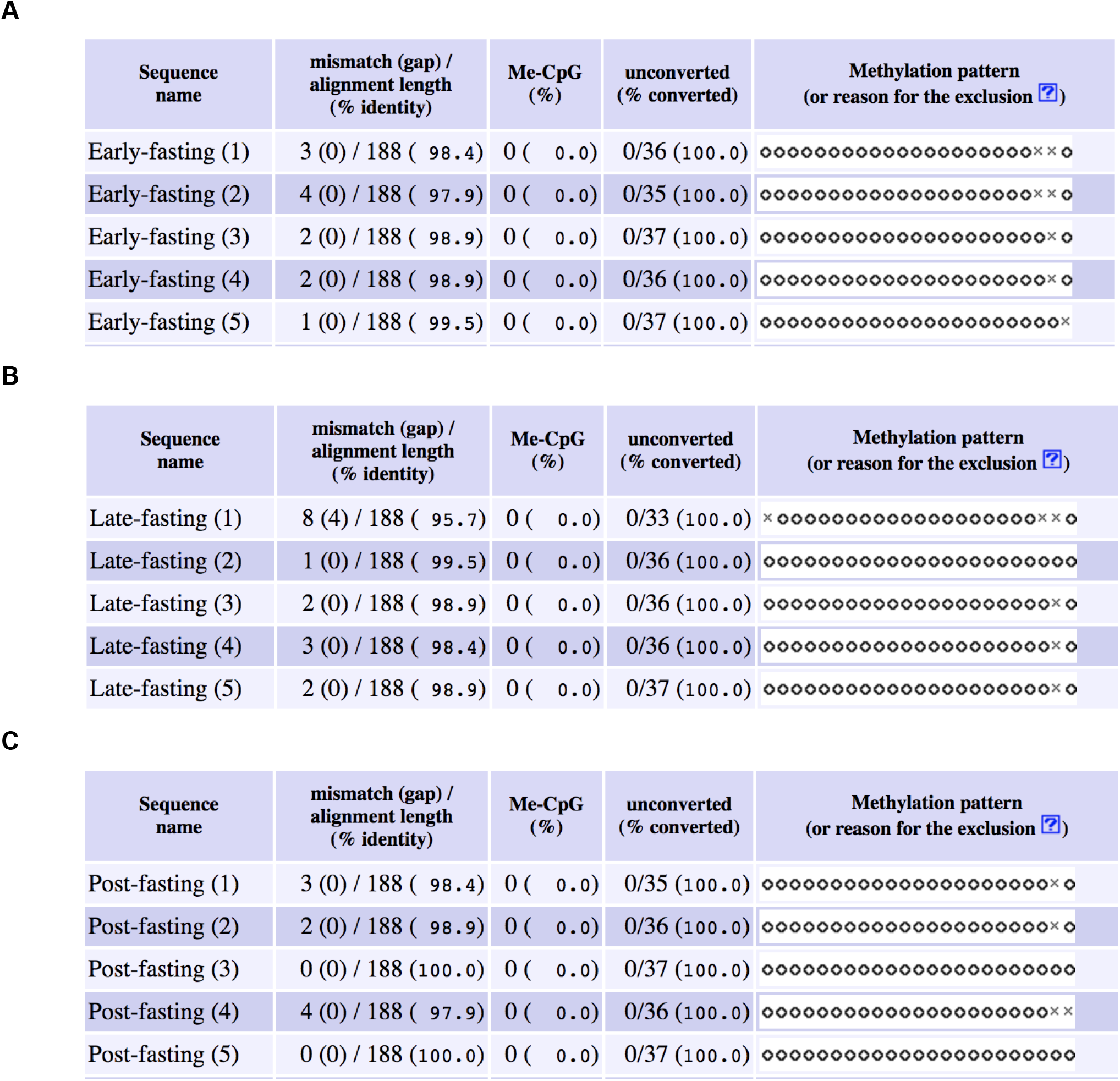
Methylation status in the GR promoter region (based on sequencing using BSP2 primers) in A) early fasting, B) late fasting, and C) post-foraging elephant seal pups. Filled-in (dark) circles indicate methylated CpG sites, and unfilled circles indicate unmethylated CpG sites. An “x” instead of a circle indicates that the CpG site was not sequenced. Data was compiled using QUMA software (http://quma.cdb.riken.jp/)

**Figure S3.**
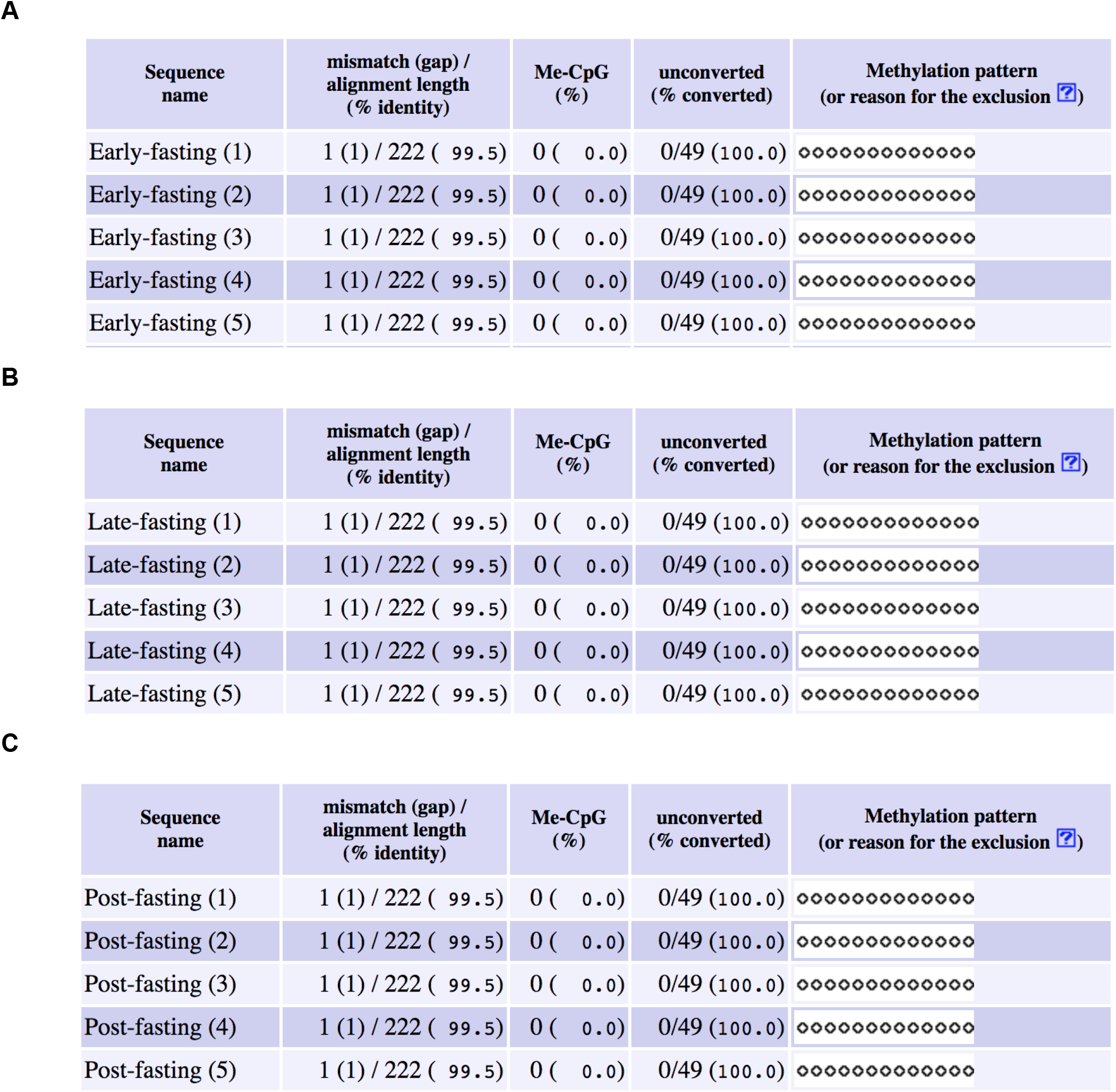
Methylation status in the GR promoter region (based on sequencing using BSP3 primers) in A) early fasting, B) late fasting, and C) post-foraging elephant seal pups. Filled-in (dark) circles indicate methylated CpG sites, and unfilled circles indicate unmethylated CpG sites. An “x” instead of a circle indicates that the CpG site was not sequenced. Data was compiled using QUMA software (http://quma.cdb.riken.jp/).

**Supplementary Table 1.** Raw methylation data.

| **Early-fasting pups** |  |  |
| --- | --- | --- |
| **Sample** | **Avg OD** | **% meth** |
| MA19-01 | 0.2765 | 0.0698683 |
| MA19-02 | 0.4395 | 0.09464128 |
| MA19-03 | 0.4215 | 0.0915221 |
| MA19-04 | 0.8245 | 0.20533107 |
| MA19-05 | 0.469 | 0.09998482 |
| MA19-06 | 1.168 | 0.3674003 |
| **Late-fasting pups** |  |  |
| **Sample** | **Avg OD** | **% meth** |
| MA19-10 | 1.046 | 0.29274625 |
| MA19-11 | 1.539 | 1.93736146 |
| MA19-12 | 1.283 | 1.8396459 |
| MA19-13 | 1.846 | 2.03505365 |
| MA19-14 | 0.677 | 1.49628588 |
| MA19-15 | 0.6345 | 1.54262287 |
| **Post-foraging** |  |  |
| **Sample** | **Avg OD** | **% meth** |
| MA19-19 | 0.3835 | 0.08527063 |
| MA19-20 | 1.0215 | 0.27969252 |
| MA19-21 | 0.8565 | 0.20571372 |
| MA19-22 | 1.0295 | 0.28388967 |
| MA19-23 | 1.0715 | 0.30698024 |
| **Dexamethasone treatment** |  |  |
| 6hr/0.1uM/1 | 0.5205 | 3.006013759 |
| 6hr/0.1uM/2 | 0.419 | 1.704477827 |
| 6hr/0.1uM/3 | 0.3905 | 1.453467278 |
| 6hr/1uM/1 | 0.444 | 1.960112122 |
| 6hr/1uM/2 | 0.397 | 1.507247403 |
| 6hr/1uM/3 | 0.463 | 2.179741432 |
| 6hr/100uM/1 | 0.369 | 1.28887927 |
| 6hr/100uM/2 | 0.328 | 1.024899691 |
| 6hr/100uM/3 | 0.462 | 2.167591288 |
| 6hr/cont/1 | 0.5525 | 3.594790616 |
| 6hr/cont/2 | 0.468 | 2.241521408 |
| 6hr/cont/3 | 0.422 | 1.73330146 |
| 12hr/0.1uM/1 | 0.502 | 2.710695225 |
| 12hr/0.1uM/2 | 0.3665 | 1.270993363 |
| 12hr/0.1uM/3 | 0.373 | 1.318021722 |
| 12hr/1uM/1 | 0.432 | 1.83294708 |
| 12hr/1uM/2 | 0.4055 | 1.580589142 |
| 12hr/1uM/3 | 0.36 | 1.225643023 |
| 12hr/100uM/1 | 0.223 | 0.569882362 |
| 12hr/100uM/2 | 0.251 | 0.666434577 |
| 12hr/100uM/3 | 0.1695 | 0.422580605 |
| 12hr/cont/1 | 0.4545 | 2.078598116 |
| 12hr/cont/2 | 0.42 | 1.71403205 |
| 12hr/cont/3 | 0.427 | 1.782428077 |
| 48hr/0.1uM/1 | 0.424 | 1.752787501 |
| 48hr/0.1uM/2 | 0.3675 | 1.278117747 |
| 48hr/0.1uM/3 | 0.2895 | 0.826455439 |
| 48hr/1uM/1 | NA | NA |
| 48hr/1uM/2 | 0.412 | 1.639072936 |
| 48hr/1uM/3 | 0.4555 | 2.09024942 |
| 48hr/100uM/1 | 0.3655 | 1.263908691 |
| 48hr/100uM/2 | 0.2495 | 0.660870168 |
| 48hr/100uM/3 | 0.1475 | 0.37368249 |
| 48hr/cont/1 | 0.498 | 2.650759712 |
| 48hr/cont/2 | 0.5925 | 4.495488522 |
| 48hr/cont/3 | 0.476 | 2.344032073 |

